# AutoFocus: A hierarchical framework to explore multi-omic disease associations spanning multiple scales of biomolecular interaction

**DOI:** 10.1101/2023.09.06.556542

**Authors:** Annalise Schweickart, Kelsey Chetnik, Richa Batra, Rima Kaddurah-Daouk, Karsten Suhre, Anna Halama, Jan Krumsiek

## Abstract

Recent advances in high-throughput measurement technologies have enabled the analysis of molecular perturbations associated with disease phenotypes at the multi-omic level. Such perturbations can range in scale from fluctuations of individual molecules to entire biological pathways. Data-driven clustering algorithms have long been used to group interactions into interpretable functional modules; however, these modules are typically constrained to a fixed size or statistical cutoff. Furthermore, modules are often analyzed independently of their broader biological context. Consequently, such clustering approaches limit the ability to explore functional module associations with disease phenotypes across multiple scales. Here, we introduce AutoFocus, a data-driven method that hierarchically organizes biomolecules and tests for phenotype enrichment at every level within the hierarchy. As a result, the method allows disease-associated modules to emerge at any scale. We evaluated this approach using two datasets: First, we explored associations of biomolecules from the multi-omic QMDiab dataset (n = 388) with the well-characterized type 2 diabetes phenotype. Secondly, we utilized the ROS/MAP Alzheimer’s disease dataset (n = 500), consisting of high-throughput measurements of brain tissue to explore modules associated with multiple Alzheimer’s Disease-related phenotypes. Our method identifies modules that are multi-omic, span multiple pathways, and vary in size. We provide an interactive tool to explore this hierarchy at different levels and probe enriched modules, empowering users to examine the full hierarchy, delve into biomolecular drivers of disease phenotype within a module, and incorporate functional annotations.

## 1 Introduction

The increasing availability of high-throughput measurement technologies has led to the generation of a large number of multi-omics datasets, providing molecular snapshots of biological systems at all -omic levels of regulation^1, 2^. Such multi-omic datasets can be explored to infer molecular interactions^3–5^, or in the context of disease, to identify perturbations for a deeper understanding of pathophysiological mechanisms^6–8^. To this end, various computational methods have been developed to cluster multi-omic biomolecules into easier-to-interpret functional modules that attempt to describe alterations caused by a disease in a biological system^5, 9–13^.

Functional modules generally consist of interacting biomolecules that are coordinated, coregulated, or otherwise involved in the same biological process^14, 15^. Grouping molecules into such functional modules can often be achieved using existing functional annotations available in large databases comprised of experimentally derived interactions^16, 17^. However, these types of annotations are constrained by research bias and are limited between -omic layers, for example those between metabolomics and transcriptomics^18, 19^. Thus, while experimentally validated annotations promise to create well-supported functional modules, the lack of exhaustive annotations in a high-throughput context is often a severe limitation. Data-driven methods that infer interactions between biomolecules directly from the data are often a compelling alternative. Such methods include k-means clustering, hierarchical clustering, network approaches, principal component analysis (PCA), or other matrix factorization approaches^12, 20–23^.

A significant challenge for these data-driven methods which statistically identify modules is determining the appropriate scale of a biological process that should be deemed a module. For instance, the catabolism of carbon units of cells can be studied at various levels, such as single-molecule level (glucose or pyruvate), pathway level (glycolysis), or functional pathway group level (central carbon metabolism, **Figure 1a**)^24^. This exemplifies the concept that functional modules are not necessarily distinct processes, and that different hierarchical levels of super- and sub-modules exist^25, 26^. In addition, previous work by our group has shown that phenotypes can impact biological system at a variety of levels; certain phenotypes, for example related to specific pathological perturbations, manifest at the level of a few molecules, while others, like sex effects, impact entire pathways or pathway groups^12, 27^.

**Figure 1.**
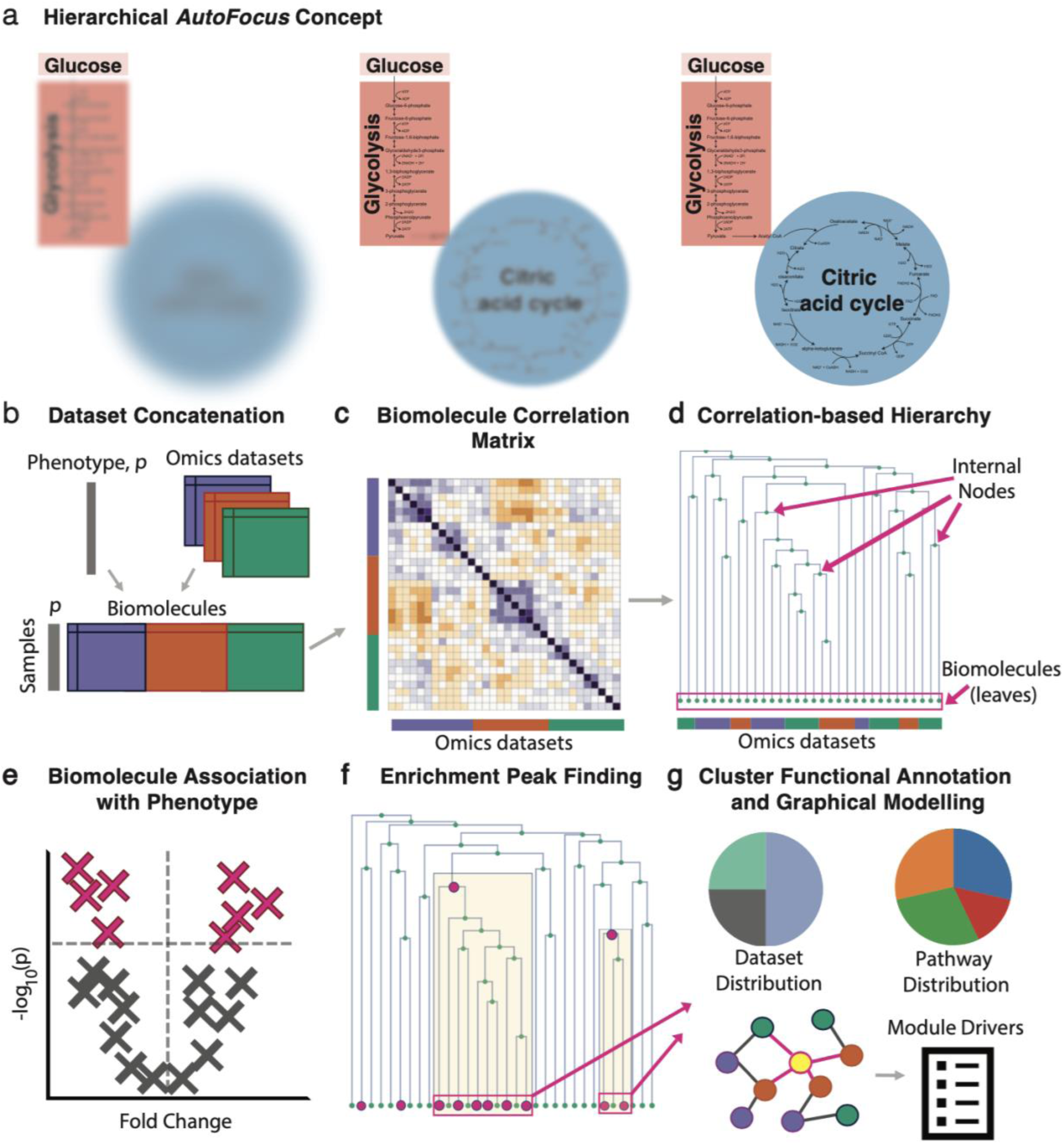
AutoFocus Method Overview. **a)** Conceptual depiction of applying “focus” to the biological process of carbon metabolism at different hierarchical levels. **b)** Multiple molecular datasets with biomolecules from the same *n* samples are concatenated into a single matrix, accompanied by sample phenotype information, *p*. **c)** Spearman correlations between molecules are calculated to generate a correlation matrix, **d)** Correlation coefficients are converted to distances to create a hierarchical tree of biomolecules, **e)** Biomolecules are univariately correlated with the phenotype of interest and filtered for statistical significance, **f)** Enrichment “peaks” are detected by performing an enrichment analysis of the “leaves” descending from each internal node, i.e., the number of significantly correlated molecules in the respective cluster. **g)** Functional annotation and module driver analysis is performed on each enriched module.

Despite the biological relevance of such hierarchies, current module identification algorithms are not designed to produce data-driven modules that can explore biological processes at multiple scales. Existing algorithms apply restrictive parameters, such as p-value cutoffs, network connectivity metrics, or desired module size, to demarcate modules at a fixed level, which are then further explored as standalone processes, disconnected from the larger biological context^10, 11, 28–30^. Thus, when analyzing the effects of a disease-phenotype on these modules, fixed-scale approaches do not reveal how a phenotype impacts different granularity levels within a module (e.g., single molecule versus pathway levels), and cannot determine how impacted modules relate to one another in the larger biological system. Further, fixed-scale modules restrict *all* phenotype associations to a single level, failing to capture the variety of scales that may exist among diverse phenotypes.

We here address the issue of identifying multi-level modules and allowing phenotype association to manifest at any scale by designing an interactive and adaptive hierarchical clustering and phenotype association approach. We introduce a new method, AutoFocus, that hierarchically structures molecular datasets, overlays phenotype association onto the hierarchy, and performs enrichment analysis to annotate functional modules within this system. The method is accompanied by an interactive application that allows a user to explore the hierarchy created by their data and provides functional insights through module annotation and the identification of module members driving phenotype association (**Figure 1**). We then apply our method to two independent datasets to validate its ability to capture known disease signal and explore new findings: Type 2 Diabetes in The Qatar Metabolomics Study on Diabetes (QMDiab, *n* = 388)^2^, which contains 12 multi-omic datasets including metabolomics, proteomics, and glycomics; and the Alzheimer’s Disease in the Religious Orders Study/Memory and Aging Project (ROS/MAP, *n* = 500)^31^, which includes a metabolomics and proteomics platform and multiple clinical phenotypes. Finally, we show how our method can easily integrate other hierarchical clustering methods into its analysis pipeline, such as the popular WGCNA clustering method^10^.

## 2 Results

### 2.1 Description of AutoFocus Framework

The AutoFocus tool enables fast clustering and phenotype association of multiple omics datasets, accompanied by an intuitive, interactive application for result exploration. Matched-sample omics datasets from any specimen, body fluid, or platform, are combined and pairwise correlated (**Figure 1b-c**). These correlations are transformed into a distance metric that is used to structure all molecules into a single dependency tree based on well-established hierarchical clustering (**Figure 1d**). Univariate associations of each molecule with a desired phenotype of interest are calculated, and significantly associated molecules, which are the “leaves” of the tree, are annotated at the bottom of the diagram (**Figure 1d-f**).

The tree is then scanned from top to bottom. For each internal node of the tree, the leaves descending from that node create a cluster (see highlighted parts of **Figure 1f**). An enrichment analysis of significant hits is performed on the molecules within that cluster. If a user-defined enrichment threshold is reached, that internal node is labeled as an “enrichment peak” (**Figure 1f**). Finally, functional annotation is performed on the modules associated with each peak along with a phenotype “driver” analysis (**Figure 1g**). Drivers are defined as module members sharing a direct, unconfounded correlation edge with the disease phenotype based on a mixed-distribution graphical model.

All AutoFocus functionalities are available as an R package at https://github.com/krumsieklab/autofocus. As input, the method accepts Excel sheets of measurements from multiple omics datasets along with dataset-specific molecular annotations and sample-specific annotations, including phenotype(s) of interest and covariate information. Accompanying the workflow in Figure 1 is an interactive Shiny application that allows a user to set an enrichment threshold and easily explore the resulting functional modules.

### 2.2 Intra- and inter-dataset relationships in the 12-dataset multi-omics QMDiab study

As all data-driven clustering methods depend on the similarity relationships between the measured variables, we first explored the correlation structure of a dataset with various omics layers to get an overview of the highly complex underlying statistical structures. The QMDiab dataset consists of 5,135 biomolecules from 8 metabolomics datasets (5 different platforms performed on plasma, 2 on urine, and 1 on saliva), 3 blood glycomics datasets, and 1 blood proteomics dataset (**Table 1**). These 12 datasets were combined and the pairwise biomolecule correlations were calculated. A systemic correlation bias was detected across the various assays: Intra-dataset correlations were systematically higher than inter-dataset correlations (**Figure 2a**). This bias persisted even in instances where the same molecule was measured on different platforms. For example, when analyzing two specific molecules, valine and leucine, measured on two almost identical plasma metabolomics platforms, we observed that valine had a higher correlation with leucine measured on the same platform than its correlation with itself measured on a different platform (**Figure 2b**). As a consequence of this bias, molecules from the same dataset tended to be in close proximity in a hierarchical structure (**Figure 2c**). This poses a problem when using correlation networks to statistically extract interactions between these molecules, a common approach to inferring biological relationships. We systematically probed the QMDiab correlation network for the optimal statistical cutoff to create a network whose edge set best models ground truth interactions. We found that this optimal cutoff differs between ground truth annotations for intra-dataset edges (metabolite pathways) and ground truth annotations for inter-dataset edges (KEGG and Recon3D-based gene-metabolite edges, **Supplementary** Figure 1)^32^. This indicates the inability of a single statistical cutoff to recover biologically relevant interactions in a multi-dataset context.

**Figure 2.**
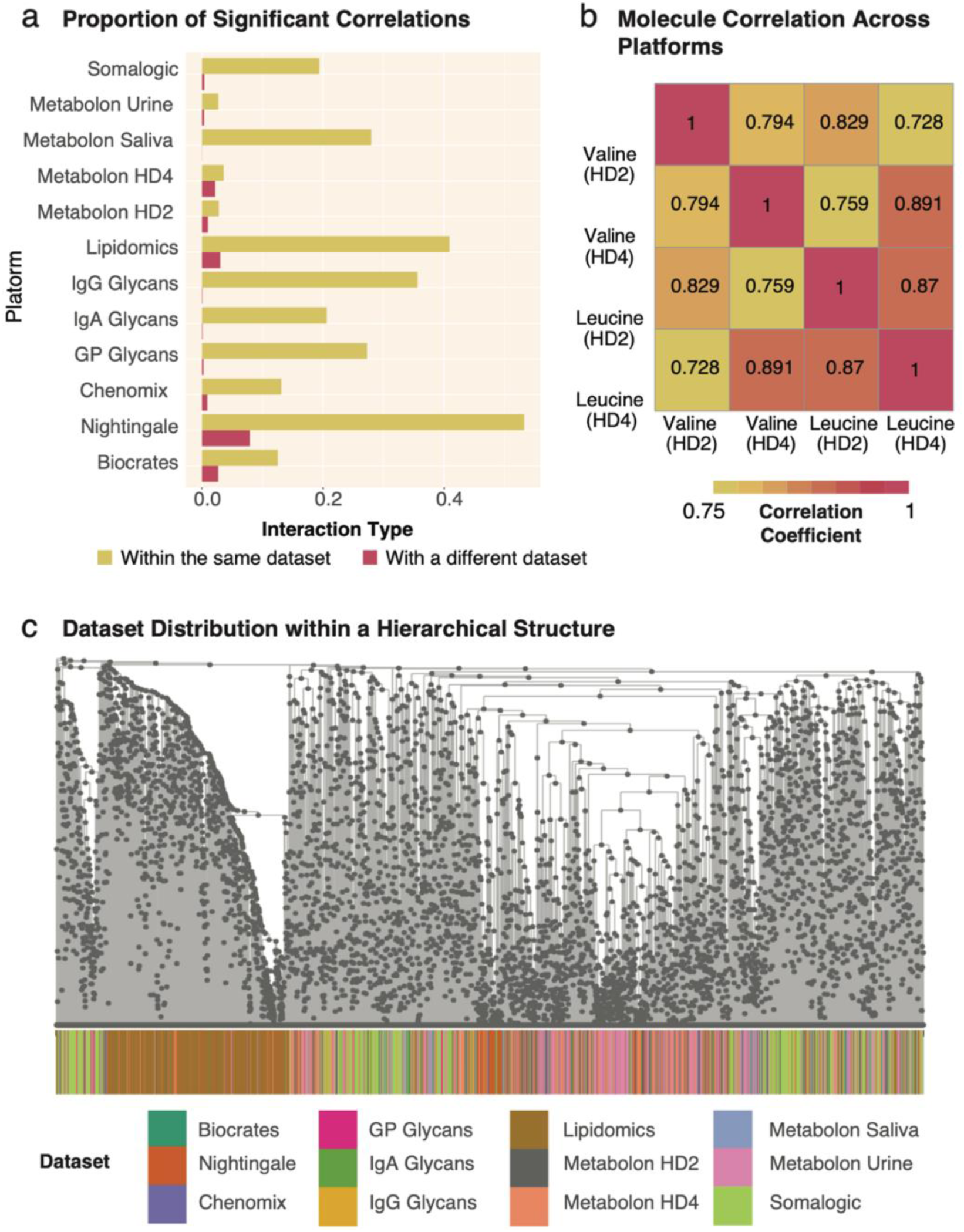
Correlation Values Within and Across Datasets. **a)** Proportion of significant correlations between biomolecules within and across datasets. For every dataset, the proportion of significant correlation coefficients within each dataset is substantially larger than across datasets. Consequently, statistical methods that depend on correlations will be biased towards intra-dataset interactions in a multiomics setting. **b)** Example correlations between two molecules measured on the sample blood samples using two similar metabolomics platforms, Metabolon Plasma HD2 and Metabolon Plasma HD4. Valine on the HD2 platform correlated stronger with Leucine measured on the same platform than with Valine on the HD4 platform. This further illustrates the tendency for stronger correlations within a dataset than between datasets. **c)** Dataset distribution in the correlation-based hierarchical structure formed on the QMDiab dataset. Strong intra-dataset correlations can be seen for lipids (brown) and to a lesser extent for proteomics (light green), as these two datasets have dense regions where they segregate from the other - omics datasets which are otherwise thought to be well integrated.

**Table 1.** Overview of -omic types, sample types, platforms, collection methods, and analyte count for the QMDiab and ROS/MAP datasets.

| Dataset | Omic Type | Sample Type | Platform | Method | # of Analytes |
| --- | --- | --- | --- | --- | --- |
| **QMDiab**<br>**(n = 410)** | Metabolomics | Plasma | Metabolon (HD2) | UHPLC/GC-MS/MS | 466 |
|  |  | Plasma | Metabolon (HD4) | UHPLC/GC-MS/MS | 843 |
|  |  | Plasma | Nightingale | NMR | 224 |
|  |  | Plasma | Biocrates p150 | FIA-MS/MS | 161 |
|  |  | Saliva | Metabolon | UHPLC/GC-MS/MS | 251 |
|  |  | Urine | Metabolon | UHPLC/GC-MS/MS | 695 |
|  |  | Urine | Chenomx | NMR | 32 |
|  | Proteomics | Plasma | SOMAscan | SOMAmer + DNA microarray | 1141 |
|  | Lipidomics | Plasma | Metabolon (Lipidyzer) | LC-MS | 1133 |
|  | Glycomics | Plasma | Genos | IgG | 39 |
|  |  | Plasma | Genos | Total N-glycans | 60 |
|  |  | Plasma | Leiden University | IgA | 90 |
| **ROS/MAP**<br>**(n= 500)** | Metabolomics | Brain Tissue | Metabolon | UHPLC/GC-MS/MS | 667 |
|  | Proteomics | Brain Tissue | ACQUITY UPLC + TSQ-Vantage MS | UHPLC-MS/MS | 7,526 |

Notably, this observation of higher correlations within the same omics layers appears to be a natural feature of multi-omics datasets. Neither AutoFocus, nor any other clustering-based method can sufficiently remove this bias, as all clustering methods rely on the associations between measured molecules. However, unlike existing methods, the hierarchical framework of AutoFocus does not apply a fixed statistical cutoff to correlations between analytes, allowing any intra- and inter-dataset relationships to naturally emerge from the data structure. As the AutoFocus method evaluates clusters at every internal node of a hierarchy, clusters formed by nodes closer to the root of the tree will encompass molecules spanning different -omics, fluids, and datasets whose relationships would have been excluded when using statistical significance-based cutoffs (**Fig 1c**). The resulting clusters are more representative of multi-omic, multi-fluidic, and multi-dataset biological interactions as compared to cutoff-dependent clustering methods.

### 2.3 AutoFocus analysis on QMDiab reveals impact of Type 2 Diabetes at multiple levels of molecular interactions

#### Systems-level Analysis of Type 2 Diabetes

The phenotype of interest used for this analysis was Type 2 Diabetes (T2D) diagnosis. After correcting for age, sex, and BMI, 188 of the 5,135 molecules were found to be significantly associated with Type 2 Diabetes (p < 0.05, Bonferroni adjusted), covering 10 of the 12 omics datasets. The IgG and IgA glycomics datasets showed no significant associations with T2D. We observed a broad distribution of signal across the hierarchical tree (**Figure 3a**), suggesting a system wide T2D effect across omics and body fluids. Certain regions of the tree had substantially denser distributions of significantly associated biomolecules, suggesting hotspots of T2D perturbation.

**Figure 3.**
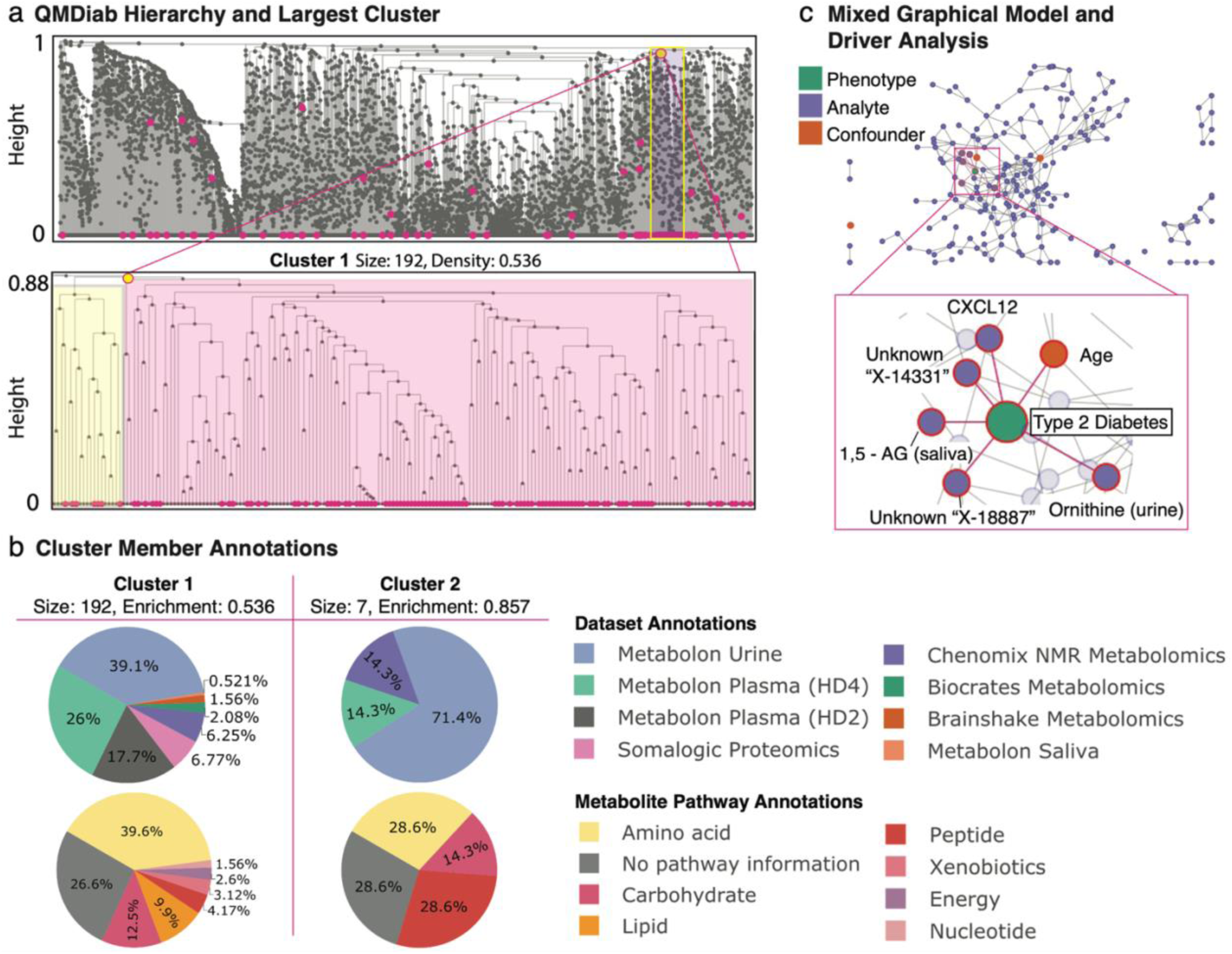
AutoFocus on the QMDiab dataset. The dataset included a total of 388 samples and 5,135 biomolecules from 12 datasets: 5 metabolomics platforms on plasma, 2 on urine, and 1 on saliva, 3 blood glycomics datasets and 1 blood proteomics dataset. **a)** View of the full hierarchical structure created from the QMDiab dataset. Magenta circles at the bottom of the tree indicate significant molecules, circles within the tree indicate modules that passed the enrichment threshold. Significant molecules were dispersed throughout the leaves of the tree and enriched modules were scattered throughout the hierarchy at a wide range of heights. The high-density region of significant molecules towards the right corresponds to the largest enriched module at the highest height. Below is a zoomed view of this module, with the left sub-tree in yellow and the right sub-tree in pink. **b)** Pie charts of the dataset and pathway makeup of the two largest modules along with their size and significant-node enrichment fraction. Pathway annotations were only available for the metabolites measured by Metabolon. **c)** Confounder-corrected mixed graphical model of the molecules in the largest module with phenotype. At the bottom, a zoomed-in view of nodes with edges to the Type 2 Diabetes phenotype which include 1,5-AG in saliva, ornithine in urine, and the CXCL12 protein, along with the confounder age and 2 unknown molecules. As these molecules are directly connected to the T2D phenotype, we mark them as statistical “drivers” of the disease in this module.

#### Type 2 Diabetes Modules

To identify T2D modules for the QMDiab dataset, we applied a “majority vote” enrichment threshold of 0.5, where at least 50% of a cluster’s members must be significantly associated with T2D for it to be designated as a T2D module. The AutoFocus method identified 21 modules, ranging in size from 2 to 192 biomolecules (**Supplementary** Figure 1 **and Supplementary Table 1**). In addition, there were 33 single-molecule modules, identified as T2D-associated molecules that did not belong to any of the 21 modules. The identified T2D modules substantially ranged in scale (**Figure 3a**), from very high correlation near the leaves at tree height 0 to low correlations near the root at tree height 1. This shows that Type 2 Diabetes manifests at various levels of the biological hierarchy, from closely connected molecules to larger pathways.

As expected, most of the smaller, highly correlated modules tended to contain molecules from only one dataset, due to the aforementioned within-dataset correlation biases, most notably within the lipidomic and proteomics datasets. However, AutoFocus identified six T2D modules that contained molecules from multiple omics or fluids (**Supplementary** Figure 1). The smaller of these modules included molecules that were measured multiple times but on different platforms, e.g., one module which was made up of pyroglutamine measured on the Metabolon HD2 and HD4 platforms. The largest module with 192 molecules (**Figure 3a**), comprising of the bulk of the T2D-associated analytes in the QMDiab dataset, brought together molecules from both metabolomic and proteomic datasets and all three body fluids in QMDiab (**Figure 3b**).

This 192-analyte module contained two sub-modules, each with substantially different functional components. The larger, right-hand “child” tree (**Figure 3a**, pink) contained molecules involved in energy metabolism, including various carbohydrates, such as mannose, glucose, and 1,5-anhydroglucitol, which are known biomarkers of diabetes^33, 34^, as well as TCA cycle metabolites like pyruvate and lactate in plasma. In addition, this module showed significant changes in the abundance of ketone bodies acetoacetate and 3-hydroxybutyrate in urine, supporting the prevalence of ketosis and ketone body secretion in T2D patients^35^.

The left sub-tree (**Figure 3a**, yellow) in this module contained biomolecules related to bone growth, mineralization, and degradation, as well as some chemokines and endothelial cell proteins. The bone degradation molecules included the proteins Osteomodulin (OMD), Integrin-binding sialoprotein (IBSP), and C-type lectin domain protein (Clec11A) and the metabolite prolylhydroxyproline in both plasma and urine^36–39^. Osteoporesis has a well-documented relationship to T2D, and although the mechanisms are not established, hypotheses for the link include inflammation and microangiopathy^40^. The presence of chemokines Stromal cell-derived factor 1 (CXCL12) and C-C motif chemokine 22 (CCL22), as well as Endothelial cell-specific molecule 1 (ESM1) in this sub-module presented potential osteoporosis links to inflammation and microangiopathy, respectively.

#### Type 2 Diabetes Module Driver Analysis

We further analyzed this module using a mixed graphical model (MGM) approach, which allowed us to differentiate direct correlations between biomolecules and T2D from indirect, statistically confounded correlations. We identified and labeled as drivers those molecules that had a direct correlation with T2D diagnosis, signified by sharing an edge in the MGM network. The MGM identified 5 biomolecules showing direct correlations with T2D, including CXCL12 and ornithine in urine, 1,5-anhydroglucitol in saliva, as well as 2 unknown urine metabolites (**Figure 3c**). The variety of drivers likely reflects the multiple functional components associated with T2D (such as hyperglycemia and inflammation).

Taken together, the AutoFocus analysis on this large T2D dataset showed the benefits of exploring multi-omics datasets with a hierarchical algorithm: First, AutoFocus was able to cluster and draw links between multiple omics and fluids into functional modules in T2D at a variety of scales within the hierarchy. Second, the granularity of the hierarchical structure allowed us to explore the functional sub-modules of an identified enrichment peak separately and in detail. For the largest QMDiab cluster associated with T2D, we were able to identify that one sub-module was enriched for energy metabolism molecules and the other for bone growth and degradation, while the peak annotations showed us how these two processes interacted together at a larger biological scale. Finally, mixed graphical models allowed us to perform a driver analysis on each module, indicating which molecules had direct statistical links to the T2D phenotype in each module, and which ones were confounded correlations.

### 2.4 AutoFocus on ROS/MAP dataset shows Alzheimer’s disease phenotype impact at different levels of the biological hierarchy

As another use case, AutoFocus was applied to an Alzheimer’s disease (AD) dataset of brain samples from the Religious Order Study (ROS) and Rush Memory and Aging Project (MAP) cohorts^31^. This dataset consisted of 8,193 biomolecules from one metabolomics and one proteomics platform, both performed on brain tissue from post-mortem samples (**Table 1**). For this analysis, we examined the association between the biomolecules and two clinical AD phenotypes simultaneously: 1) Neurofibrillary tangles (NFT), defined by the immunohistochemistry-based overall paired helical filament tau tangles load from post-mortem pathology, and 2) cognitive decline (CD), defined by the rate of change in global cognition over lifetime. These phenotypes were chosen because they represent two distinct effects of AD, molecular and cognitive.

#### Systems-level Analysis of AD phenotypes

Of the 8,193 molecules in the ROS/MAP dataset, 887 molecules significantly associated with NFT and 763 molecules significantly associated with CD (p < 0.05, adjusted p-values). All statistical models were corrected for age at death, sex, BMI, post-mortem interval, years of education, and APOE genotype. To maintain consistency with previous studies published on the ROS/MAP dataset^41^, the FDR p-value correction method was used instead of Bonferroni, leading to a dense distribution of significant hits across the tree (**Figure 4a**). Both phenotypes had robust metabolic associations, as metabolites made up 20% and 26% of significant hits in NFT and CD, respectively, even though metabolites only made up 8.14% of the underlying dataset. There were 358 overlapping molecules significantly associated with both phenotypes.

**Figure 4.**
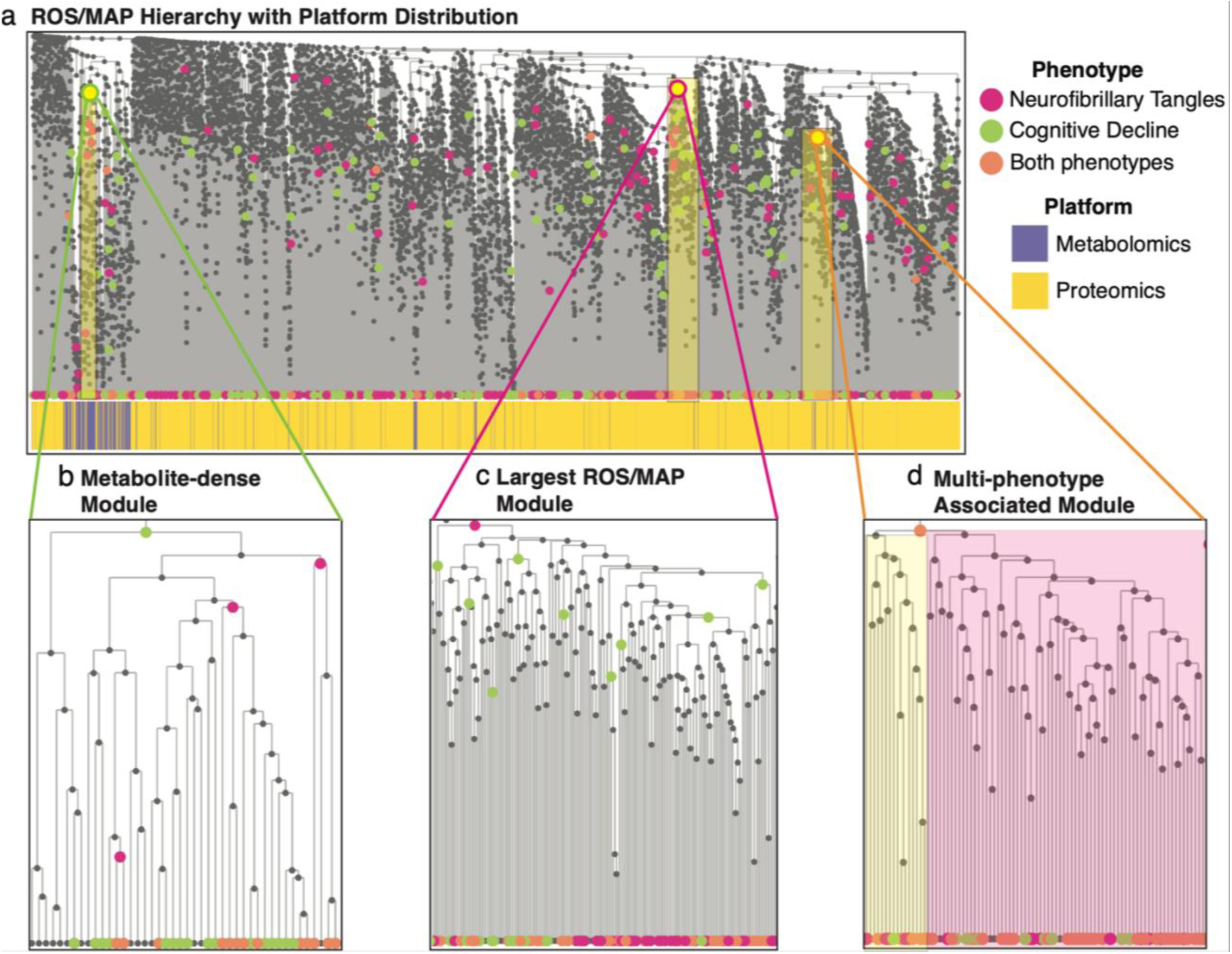
Results of running the AutoFocus method on the ROS/MAP dataset. The dataset included a total of 500 samples, which contained 8,193 biomolecules from a metabolomics platform a proteomics platform performed on post-mortem brain tissue. **a)** View of the full hierarchical structure created from the ROS/MAP dataset with two phenotypes annotated and dataset distribution below. Magenta circles represent the neurofibrillary tangles phenotype, green circles represent cognitive decline, and orange circles are overlaps between the two. Significant molecules are dispersed densely throughout the tree and enriched modules are scattered throughout the hierarchy at a large range of heights. **b)** Zoomed-in view of a metabolomics module enriched for significant hits associated with cognitive decline. This module contained metabolites related to oxidative stress and lipid peroxidation. **c)** Zoomed in view of the largest module found in the dataset which was enriched for metabolites and proteins significantly associated with neurofibrillary tangles. **d)** Zoomed-in view of the largest module enriched for both phenotypes with the left sub-tree (yellow) enriched for mitochondrial proteins and the right sub-tree (pink) enriched for proteins related to synaptic vesicle exocytosis and inhibitory neurotransmission.

#### AD Modules

A total of 170 modules were identified with a “majority vote” enrichment threshold of 0.5, with 81 modules unique to the NFT phenotype, 82 unique to the CD phenotype, and 7 modules associated with both phenotypes. There were 462 single-molecule modules that did not belong to any of the 170 modules. The multi-molecule modules ranged in size from 2 to 165 biomolecules (**Supplementary** Figure 2**, Supplementary Table 2**). Similar to the QMDiab dataset, modules associated with both phenotypes ranged drastically in tree height across the tree, from 0 to 0.85 (**Figure 4a**).

An interesting feature arising from applying AutoFocus on two phenotypes was the nesting of enrichment peaks, where the peak of one phenotype was a descendant of a peak of the other. As the NFT and CD phenotypes had a large overlap of significantly associated molecules, their enriched modules tended to occupy similar regions of the tree. Despite this considerable overlap, only 7 internal nodes were identified as enrichment peaks for both phenotypes (**Figure 4a**, orange nodes). For all other regions of the hierarchy where both phenotypes had overlapping significant hits, modules enriched for one phenotype contained descendent sub-modules enriched for the other phenotype (**Figure 4b-c**). This nesting highlights how different phenotypes within a single disease can manifest at different scales of biological processes, where cognitive decline may be associated with a biological process at a higher level than neurofibrillary tangles pathology, and vice versa.

The ROS/MAP hierarchical structure was strongly affected by the dataset correlation bias as metabolites were condensed within the tree, leading to only 6 of the 170 modules being multi-omic (**Figure 4a, Supplementary Table 2**). Within the dense metabolomic region of the ROS/MAP hierarchy, CD had more significant metabolite associations, which resulted in a higher enrichment peak (larger cluster) than the NFT phenotype. This 44-metabolite module was enriched for antioxidants and lipid peroxidation metabolites^42, 43^, indicating that CD interacts with oxidative stress metabolism at a higher biological scale than NFT (**Figure 4b, Supplementary Table 2**).

Of the 6 multi-omic modules was the largest module in the tree, which significantly associated with the NFT phenotype (**Figure 4c**). This module contained proteins and metabolites involved in a variety of processes; one sub-module showed multi-omic dysregulation of arginine flux, degradation, and metabolism^44–47^, one sub-module contained proteins associated with inflammatory mediator TNF-⍺^48–50^, while an adjacent sub-module contained glycosylation proteins^51^. In contrast, the CD phenotype had enrichment peaks for the NFT sub-modules involved in arginine metabolism and inflammation, but not for the region associated with protein glycosylation. This indicates that protein glycosylation has an NFT-specific association, and thus the NFT phenotype is associated at a higher level in the biological hierarchy for this process than the CD phenotype.

One of the 7 modules identified for both phenotypes was a proteomic module combining two functionally distinct sub-modules. The left child (**Figure 4d**, yellow) contained proteins largely involved in mitochondrial processes including mitochondrial membrane trafficking ^52, 53^ and mitochondrial gene expression^54^. The right child (**Figure 4d**, pink) was enriched for proteins related to synaptic vesicle exocytosis and inhibitory neurotransmission ^55–59^. As this was a module where both phenotypes were enriched at the same level, these processes do not seem to be specific to either phenotype.

In summary, overlaying these two phenotypes on the ROS/MAP hierarchy demonstrated the difference in biological manifestation of cognitive decline and tau neurofibrillary tangles in Alzheimer’s disease. These differences highlight phenotype-specific processes, while modules equally enriched for both phenotypes indicate more universal disease processes that may not be attributable to a single phenotype.

### 2.5 Integration with WGCNA Clustering

While the AutoFocus tool builds hierarchical trees using standard agglomerative hierarchical clustering from pairwise correlations (see **Methods 3.3.1**), the framework is applicable to any input tree structure. For example, one of the most widely used clustering methods for omics data, weighted gene co-expression network analysis (WGCNA), derives a hierarchical structure using a “topological overlap matrix” (TOM) of a co-expression network^10^. WGCNA trees can be directly integrated into the AutoFocus framework.

To investigate how a clustering method can affect AutoFocus cluster analysis, the QMDiab dataset was rerun using WGCNA’s TOM-based hierarchical structure (**Supplementary** Figure 3). Briefly, while the TOM-based hierarchy produced a very different cluster structure from the correlation-based clustering method, AutoFocus was still able to identify widespread T2D perturbations of energy metabolism with the new hierarchy, similar to those found in the initial analysis. The dataset correlation bias persisted in the WGCNA hierarchy, although while this manifested the most strikingly in the lipidomics dataset in the correlation-based hierarchy (**Fig 2c**), the proteomics dataset was highly segregated in the TOM-based tree (**Supplementary** Figure 2a). As the bone degradation pathway identified in the initial analysis was highly driven by dysregulation of proteins, this process was not recaptured in the TOM hierarchy. Overall, the TOM-based hierarchy was able to recapture only a portion of the key T2D-affected processes identified by the correlation-based hierarchy.

## 3 Discussion

The AutoFocus method provides a novel computational approach for identifying disease-perturbed, multi-omic modules of biomolecules at various resolutions of biological hierarchy. By testing for enrichment at each internal node in a hierarchical tree, AutoFocus allows relationships between molecules across all platforms, fluids, and omics to be analyzed in the context of phenotypic perturbations. The identified modules are better able to model multi-omic, multi-fluidic, and multi-dataset biological interactions as compared to clustering methods which rely on modules defined at a fixed level and explored as standalone processes. The hierarchical framework allows for the exploration of one or more phenotypes at fine granularity or at a larger, zoomed-out scale. Furthermore, the method’s implementation in an interactive application makes navigation of the complex biological structure, and the modules within, easy and intuitive.

We applied AutoFocus to two multi-omic datasets, QMDiab and ROS/MAP. For both datasets, AutoFocus was able to find a multitude of disease-associated modules at various levels of correlation. For the type 2 diabetes (T2D) phenotype in QMDiab, AutoFocus was able to detect multi-omic modules enriched for known T2D associated processes, such as energy metabolism pathways and bone degradation, distinguishing them as separate but related processes. We were able to integrate the TOM-based hierarchical structure of the WGCNA method into AutoFocus to identify discrepancies in the module results stemming from the underlying hierarchy. Applying AutoFocus to the ROS/MAP Alzheimer’s disease dataset with multiple phenotypes, we were able to distinguish the different scales at which two different pathophenotypes associated with dysregulated processes within a single disease. Without the hierarchical perspective and tool allowing us to explore multiple levels within our dataset, neither of these findings would have been possible.

The main limitation of the AutoFocus method is the multi-omic dataset correlation bias in which intra-dataset correlations are systematically higher than inter-dataset correlations. This bias affects the hierarchical structure between the molecules, and therefore the modules identified by the AutoFocus algorithm will be more likely to contain relationships within one dataset than cross-dataset interactions. Notably however, this bias will affect any method that uses statistical similarity measures between molecules. By testing clusters at all levels of the hierarchy rather than cutting clusters into disparate groups that potentially sever ties between datasets, the AutoFocus design increases the likelihood of identifying multi-omic modules if they exist.

In conclusion, AutoFocus is a new approach to detect modules in complex, multi-omics data at any scale of association. It allows for multiple phenotype comparison and comes with an interactive Shiny app for result exploration. Our results show that AutoFocus is effective at identifying interactions between biological systems and disease perturbations and can distinguish molecular modules affected by different phenotypes in complex disease.

## 4 Methods

### 4.1 Datasets

The QMDiab study was conducted at the Dermatology Department of Hamad Medical Corporation (HMC) in Doha, Qatar. The study population was predominantly of Arab, South Asian, and Filipino descent, with participants falling between the ages of 23 and 71. Data was collected between February and June of 2012; collection and sampling methods have been previously described elsewhere^60^. The study was approved by the Institutional Review Boards of HMC and Weill Cornell Medicine-Qatar (WCM-Q). Written informed consent was obtained from all participants. For the analysis described in this paper, we included data from 388 subjects (192 females, 196 males; 195 diabetic, 193 non-diabetic).

The Religious Order Study (ROS) and Rush Memory and Aging Project (MAP) are two studies conducted by the Rush Alzheimer’s Disease Center. ROS started recruiting individuals from religious communities across the United States in 1994, and MAP started recruiting individuals from a wide range of backgrounds and socio-economic statuses from Northeastern Illinois in 1997. Data collection and sampling methods have been previously described elsewhere^41^. For this study, data from post-mortem tissue of 500 subjects was included (352 females, 148 males; 220 with Alzheimer’s Disease, 119 with mild cognitive impairment, 153 with no cognitive impairment, 8 with other forms of dementia). Both cohorts were approved by an institutional review board of Rush University Medical center. All participants provided informed consent, an Anatomic Gift Act, and a repository consent to allow their data and biospecimens to be shared.

### 4.2 Multi-omic measurements

#### 4.2.1 QMDiab

Plasma metabolomic profiling was performed by running plasma samples through 5 separate platforms: 1) The Metabolon Inc. HD2 platform, which uses non-targeted ultrahigh-performance liquid chromatography (UHPLC) and gas chromatography (GC) separation coupled with mass spectrometry (MS/MS)^61^. This yielded 466 measured metabolites. 2) The Metabolon UHPLC-MS/MS and GC-MS/MS HD4 platform (843 metabolites). 3) The Metabolon Lipidyzer^TM^ platform, which resolved fatty acid side chains (1,133 lipids)^62^. 4) The Biocrates Life Sciences AG AbsoluteIDQ^TM^ p150 metabolomics kit, which used targeted flow injection analysis tandem mass spectrometry (FIA-MS/MS) from (161 molecules)^63^. 5) The targeted Nuclear Magnetic Resonance (NMR) platform of Nightingale Ltd. (224 metabolites)^64^.

Urine metabolomic profiling was performed through non-targeted ultrahigh-performance liquid chromatography and gas chromatography separation, coupled with mass spectrometry on the Metabolon Inc. HD2 platform (695 metabolites) and the targeted proton Nuclear Magnetic Resonance (^1^H NMR) platform of Chenomx, Inc. (32 metabolites)^65^.

Saliva metabolomic profiling was performed through non-targeted ultrahigh-performance liquid chromatography and gas chromatography separation, coupled with mass spectrometry on the Metabolon Inc. platform (251 metabolites).

Glycomics profiling was performed on 3 separate platforms; 356 plasma samples were sent to Genos, Ltd. (Zagreb, Croatia) for the analysis of total plasma N-glycosylation using ultra-performance liquid chromatography (UPLC) and IgG Fc N-glycosylation using liquid chromatography mass spectrometry 18lycol-profiling^66^ (39 and 60 measured glycans, respectively). IgA glycomics measurements were collected at Leiden University Medical Center using UPLC coupled to a quadrupole-TOF-MS, resulting in 90 measured IgA molecules as previously described (Dotz et al., 2021, Momcilovic et al,2020).

Plasma proteomics profiling was performed on 356 samples at the WCM-Q proteomics core, using the SOMAscan assay (version 1.1) protocols and instrumentation provided and certified by SomaLogic Inc. (Boulder, CO)^67^ (1,141 proteins).

#### 4.2.2 ROS/MAP

For 500 of the brain tissue samples of the ROS/MAP cohort, brain metabolomic profiling was performed through non-targeted ultrahigh-performance liquid chromatography and gas chromatography separation, coupled with mass spectrometry on the Metabolon Inc. platform (667 metabolites)^41^. Brain proteomic profiles were collected on 265 ROS/MAP samples using tandem mass tag (TMT)-MS and downloaded from the AMP-AD Knowledge Portal (https://adknowledgeportal.synapse.org, 7,526 proteins), details of data collection and processing have been previously described^68^.

### 4.3 Data preprocessing

#### QMDiab

For each dataset, samples with more than 20% missing molecules and molecules with more than 10% missing samples were removed. Molecular abundance levels were then probabilistic quotient normalized to correct for sample-wise variation^69^ and log-transformed. Data was then scaled, and all outliers with abundance levels above 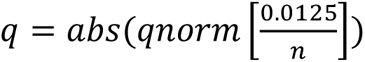, with *n* representing the number of samples, were set to missing.

Missing values were imputed using a k-nearest neighbors (k-nn) imputation method^70^. All data preprocessing was performed with the maplet R package^71^.

#### ROS/MAP

The ROS/MAP preprocessing steps have been outlined in Batra et al.^41^ and Johnson et al^68^. Briefly, metabolites with over 25% missing values were filtered out, samples were quotient-normalized and subsequently log-transformed. Outlier samples were removed using the local outlier factor method and abundance level outliers were set to NA. Missing values were imputed with a k-nn algorithm. Proteins were log_2_-transformed and corrected for batch effects using ‘median polish’ approach. Missing values and outliers were treated with same approach as the metabolomics data. Duplicated proteins with same Uniprot IDs were averaged.

After pre-processing the individual dataset, the data matrices were concatenated into a final data matrix. For the QMDiab dataset, this final data matrix consisted of 388 samples and 5,135 analytes, and for the ROS/MAP dataset, this final data matrix consisted of 500 samples and 8,193 analytes.

### 4.4 AutoFocus Method

#### Hierarchical Clustering

Once all datasets were preprocessed and concatenated (**Figure 1a**), the data matrix was hierarchically clustered. The distance metric between two analytes was derived as one minus the absolute value of their Spearman correlation value such that the stronger the correlation (either positive or negative), the closer the analytes were in the hierarchy. This distance matrix was transformed into a hierarchical structure using the average-linkage method, which has been shown to maximize the cophenetic correlation between a hierarchical structure and its correlation-based distance matrix as compared to other common linkage methods^72^ (**Figure 1c**). On the hierarchical tree, “leaf” nodes represented biomolecules. Internal nodes represented the root of all their leaf descendants; therefore, a cluster was defined at each internal node. Each internal node also had a right and left child, which could be either a leaf or another internal node.

#### Univariate Analysis

All measured molecules were associated with a phenotype of interest, *p*, using a linear model with added confounding terms to correct for applicable covariates (e.g., age, sex, BMI). P-values from this linear model were used to determine molecule significance after adjustment for multiple hypothesis testing.

#### Enrichment “Peak” Calculation

To find phenotype association enrichment among clusters of the hierarchical tree, the internal nodes of the hierarchy were scanned from top to bottom. At each internal node, the set of leaves descending from that internal node was considered; if the proportion of these leaves that were significantly associated with the phenotype of interest surpassed a user-defined enrichment threshold, this internal node was labeled as an enrichment “peak”. Once a cluster was found at which this enrichment point was met, the scanning stopped for its descendants as we reached the highest level at which disease signal was detected at the desired enrichment threshold.

This process sometimes resulted in “piggy-backers”, defined as peaks that reached the enrichment only due to one child reaching the enrichment threshold, and the joining of the two children diluted the signal (reduced the fraction of significant molecules in the cluster). Once all peaks had been identified in the hierarchy, each peak was assessed for the individual contributions from either child. A peak whose signal could be attributed to a single child was removed (details in **Supplementary** Figure 4), the child node that meets enrichment was labeled as a peak instead, and the other child that did not meet the threshold continued to be scanned. This iterative process continued until all piggy-backers were removed (**Supplementary** Figure 4).

#### Cluster Driver Analysis

Once enrichment peaks were identified, an additional analysis was performed on molecules within the biological cluster descending from each peak to identify potential drivers of the disease signal. While a significant univariate association indicated a biological link between a molecule and a phenotype, this effect could have been indirect, meaning the association was relayed through an intermediate variable that was directly associated with the phenotype. Therefore, a driver analysis was performed to identify which molecules had a direct effect.

To this end, the data matrix consisting of abundance data from the molecules descending from the enriched peak was combined with the phenotype vector and all covariates and used to build a mixed graphical model using the **mgm** package in R^73^. Graphical models use conditional dependency estimates between molecules, covariates, and disease diagnosis to extract direct correlations and to exclude indirect effects through confounding. Mixed graphical models in particular are capable of generating the conditional independence structure of many underlying distributions, including Gaussian, Poisson, and categorical ^74^.

For our application, molecules and/or covariates were labeled as drivers of a disease phenotype if they shared an edge with the disease phenotype in the resulting MGM graph, as they shared a direct correlation with the phenotype.

#### AutoFocus code and interactive tool

The AutoFocus method is accompanied by an interface developed using the Shiny app environment^75^ under R version 4.2.2. The code for the app is freely available as a GitHub at https://github.com/krumsieklab/autofocus.

### 4.5 Runtime performance

Running the AutoFocus method can be parallelized over multiple CPUs to reduce computation time for large datasets. Generating the results of the ROS/MAP dataset with 8,193 molecules and two phenotypes took 4.5 hours on a single 2.7 GHz Quad-Core Intel Core i7 CPU, with most of this time spent creating the MGMs for each cluster.

### Data availability

The preprocessed, concatenated QMDiab dataset used in this paper can be found at https://doi.org/10.6084/m9.figshare.23934933.v1.

The ROS/MAP data used in this paper can be obtained from two sources: (1) Metabolomics and proteomics data for the ROS/MAP cohort are available via the AD Knowledge Portal (https://adknowledgeportal.org). The AD Knowledge Portal is a platform for accessing data, analyses, and tools generated by the Accelerating Medicines Partnership (AMP-AD) Target Discovery Program and other National Institute on Aging (NIA)-supported programs to enable open-science practices and accelerate translational learning. The data, analyses, and tools are shared early in the research cycle without a publication embargo on secondary use. Data is available for general research use according to the following requirements for data access and data attribution (https://adknowledgeportal.org/DataAccess/Instructions). For access to content described in this manuscript see: http://doi.org/10.7303/syn26401311. (2) The full complement of clinical and demographic data for the ROS/MAP cohort are available via the Rush AD Center Resource Sharing Hub and can be requested at https://www.radc.rush.edu.

### Code availability

Codes used in this study are available at the GitHub repository https://github.com/krumsieklab/autofocus.

## Author Contributions

K. S. and A. H. provided the QMDiab data. R. B. and R.K. provided preprocessed ROS/MAP data. A. S. and J. K conceived of and designed the research study. A. S. preprocessed the QMDiab dataset, wrote the main functionality of the AutoFocus algorithm and shiny application, analyzed all data and results. K. C. formalized all code, added key shiny functionality, and made the AutoFocus R package. A. S., R.B., and J. K. wrote the manuscript. All authors gave final approval to publish.

## Acknowledgements

This study was supported by Biomedical Research Programme funds at Weill Cornell Medicine in Qatar, a program funded by the Qatar Foundation. The funders had no role in the study design, data collection and analysis, decision to publish or preparation of the manuscript.

The results published here are in whole or in part based on data obtained from the AD Knowledge Portal. Study data were provided by the Rush Alzheimer’s Disease Center, Rush University Medical Center, Chicago. Additional phenotypic data can be requested at www.radc.rush.edu.

ROSMAP metabolomics data is provided by the Alzheimer’s Disease Metabolomics Consortium (ADMC). The investigators within the ADMC, not listed specifically in this publication’s author’s list, provided data along with its pre-processing and prepared it for analysis, but did not participate in analysis or writing of this manuscript. A complete listing of ADMC investigators can be found at: https://sites.duke.edu/adnimetab/team/. The Metabolon datasets were generated at Metabolon and pre-processed by the ADMC.

