## Supplemental Figures 1-5 for "AutoFocus: A hierarchical framework to explore multi-omic disease associations spanning multiple scales of biomolecular interaction"

**Supplementary Materials**

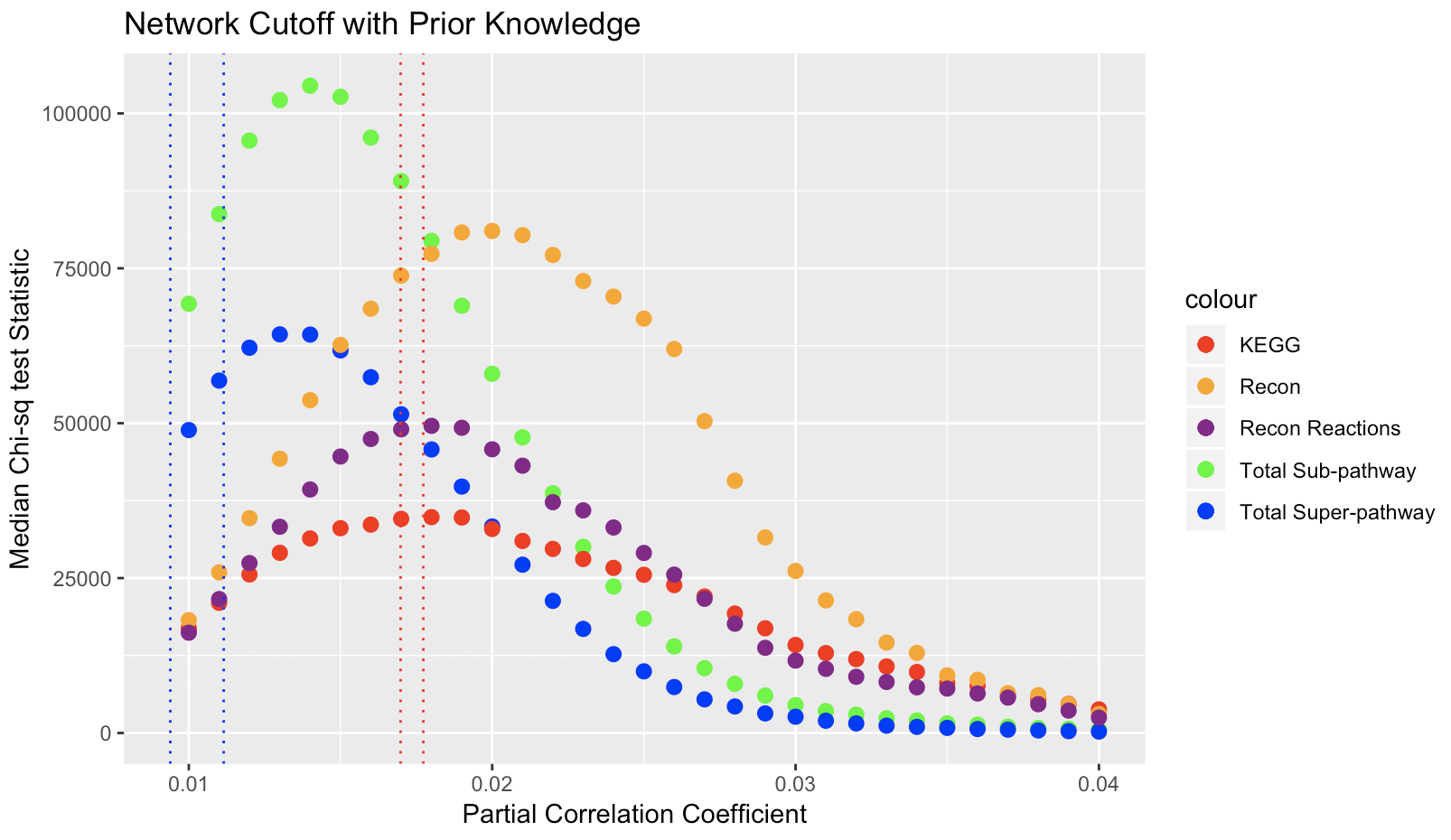

### Supplementary Figure 1 - Optimum network cutoff search for QMDiab data

**Supplementary Figure 1.** **Prior knowledge-informed optimum network cutoff search for QMDiab data.** Cutoff optimization graph for QMDiab dataset using annotations from the KEGG database (red) and Recon 3D database (molecular annotations in orange, reaction annotations in purple), as well as the sub- (green) and super- (blue) pathway annotations of the Metabolon platform. Bonferroni-adjusted p-value cutoffs are demarcated in blue vertical dashed lines at p = 0.01 (left) and p = 0.05 (right), while FDR-adjusted cutoffs are similarly demarcated in red.

Using the method outlined in Benedetti et al^1^, biomolecules of the QMDiab dataset were annotated using annotations from the KEGG^2^ and Recon 3D^3^ databases, as well as sub- and super- pathway annotations that describe biochemical processes and broader metabolite groups, respectively, for molecules measured on Metabolon platforms. Briefly, we created a Gaussian graphical model (GGM) of the QMDiab biomolecules and set the network cutoff at a partial correlation coefficient ranging from 0.01 to 0.04 (a partial correlation value greater than the cutoff would indicate an edge present in the network). For each annotation set, we performed a Chi-square test on an overlap between the GGM and the known pathways. Specifically, we calculated a contingency table which classified pairs of biomolecules based on whether an edge between them appears in the GGM and they share the same annotation (true positive), they only share a GGM edge (false positive), they only share an annotation (false negative), or they share neither an edge nor an annotation (true negative). The optimal cutoff is defined as that with the highest test statistic. For the QMDiab dataset, the optimal cutoff varied per annotation set, with the omic specific annotations (sub- and super- pathways) being optimal at a more stringent cutoff, while the cross-omic annotation sets (KEGG and Recon) being optimal at a higher coefficient. This lack of agreement illustrates the correlation-based platform bias within the QMDiab dataset, as there does not exist a single optimal correlation cutoff.

Supplementary Figure 2 - QMDiab Enriched Module Composition
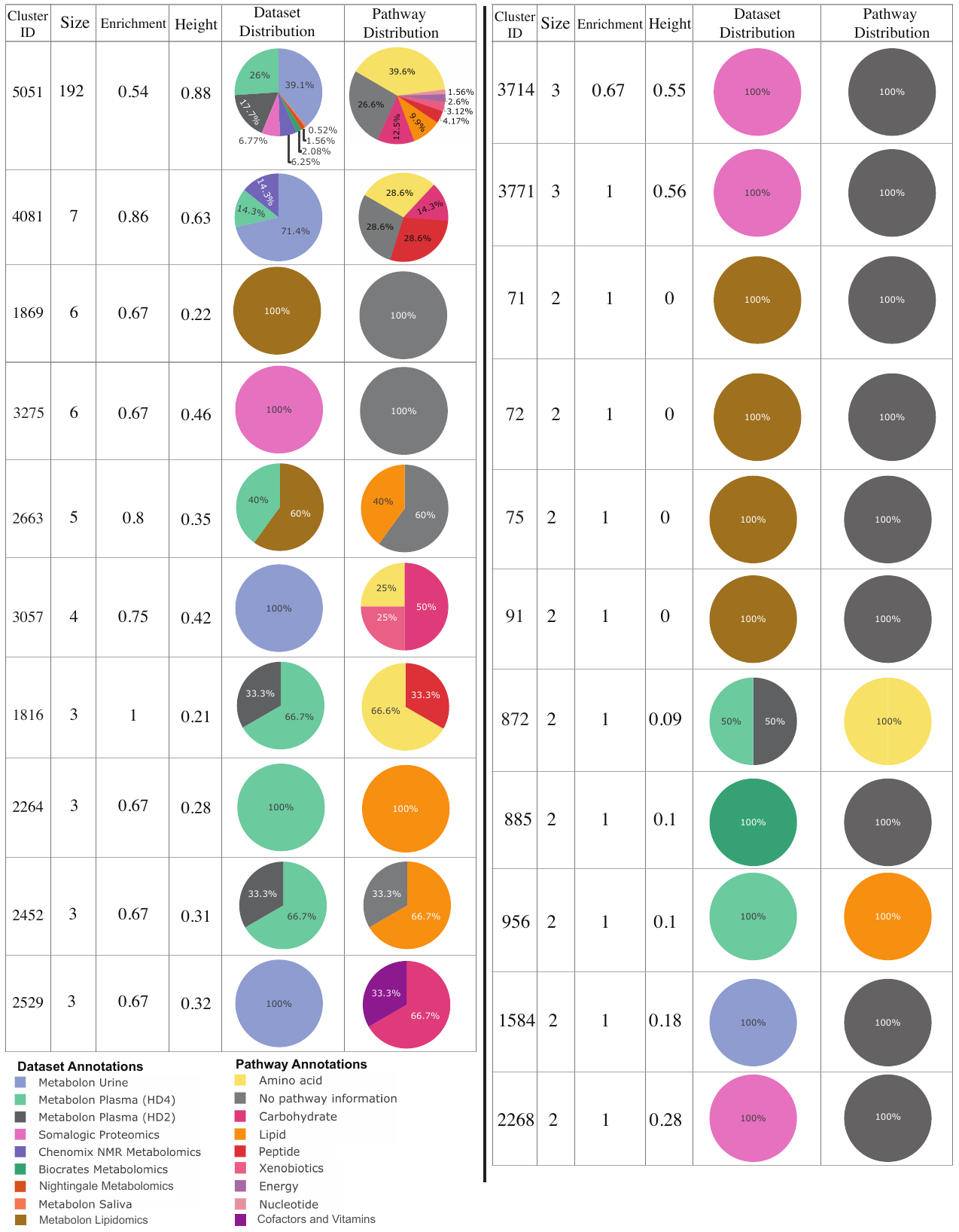

**Supplementary Figure 2.** Table of QMDiab module information. Listed are the modules’ ID (for cross referencing with **Supplementary Table 1**), size (number of molecules), enrichment (proportion of significant phenotype-associated molecules), height on the hierarchy, and distributions of the members’ datasets and pathways (Metabolon platforms only).

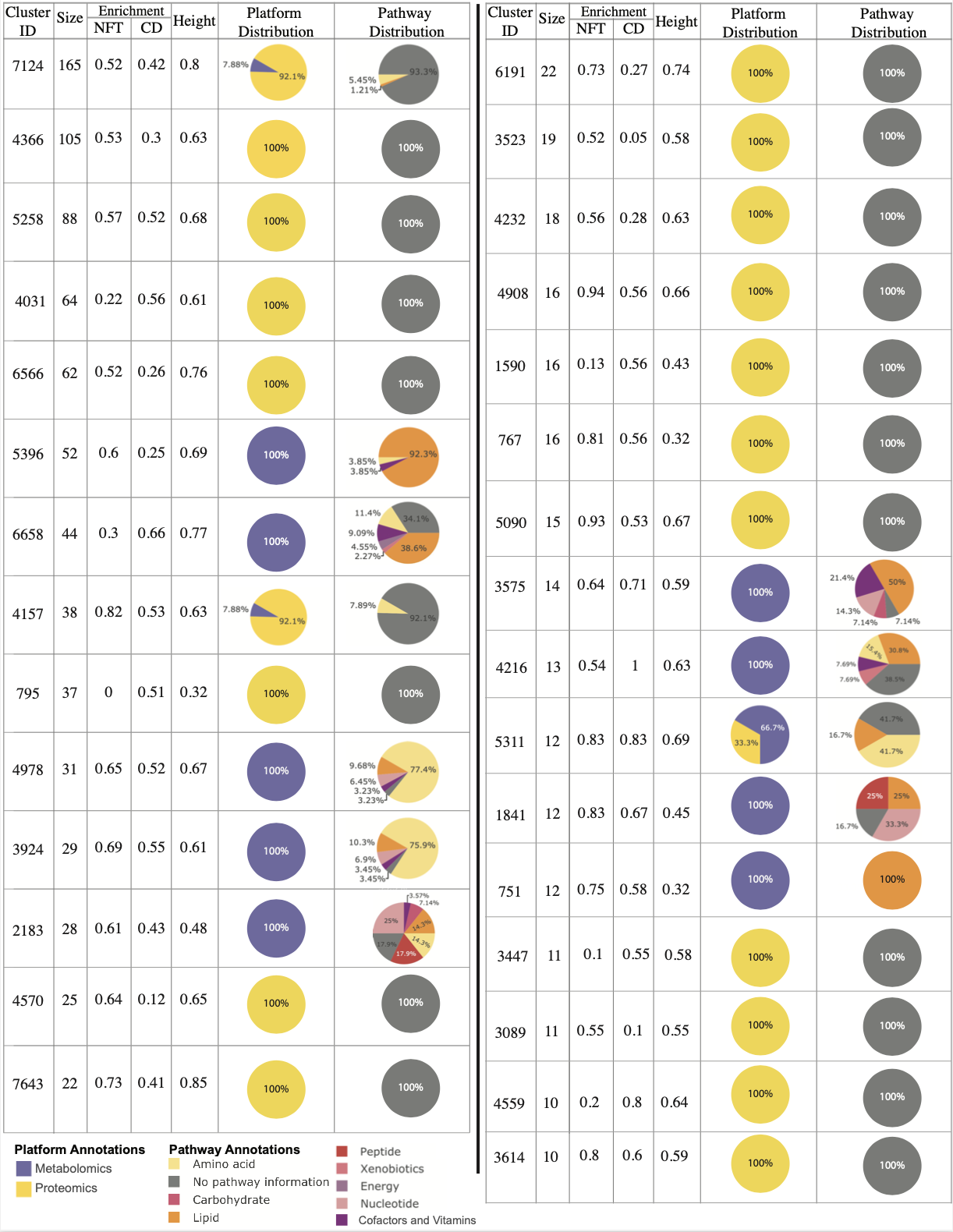
Supplementary Figure 3 – ROS/MAP Enriched Module Composition

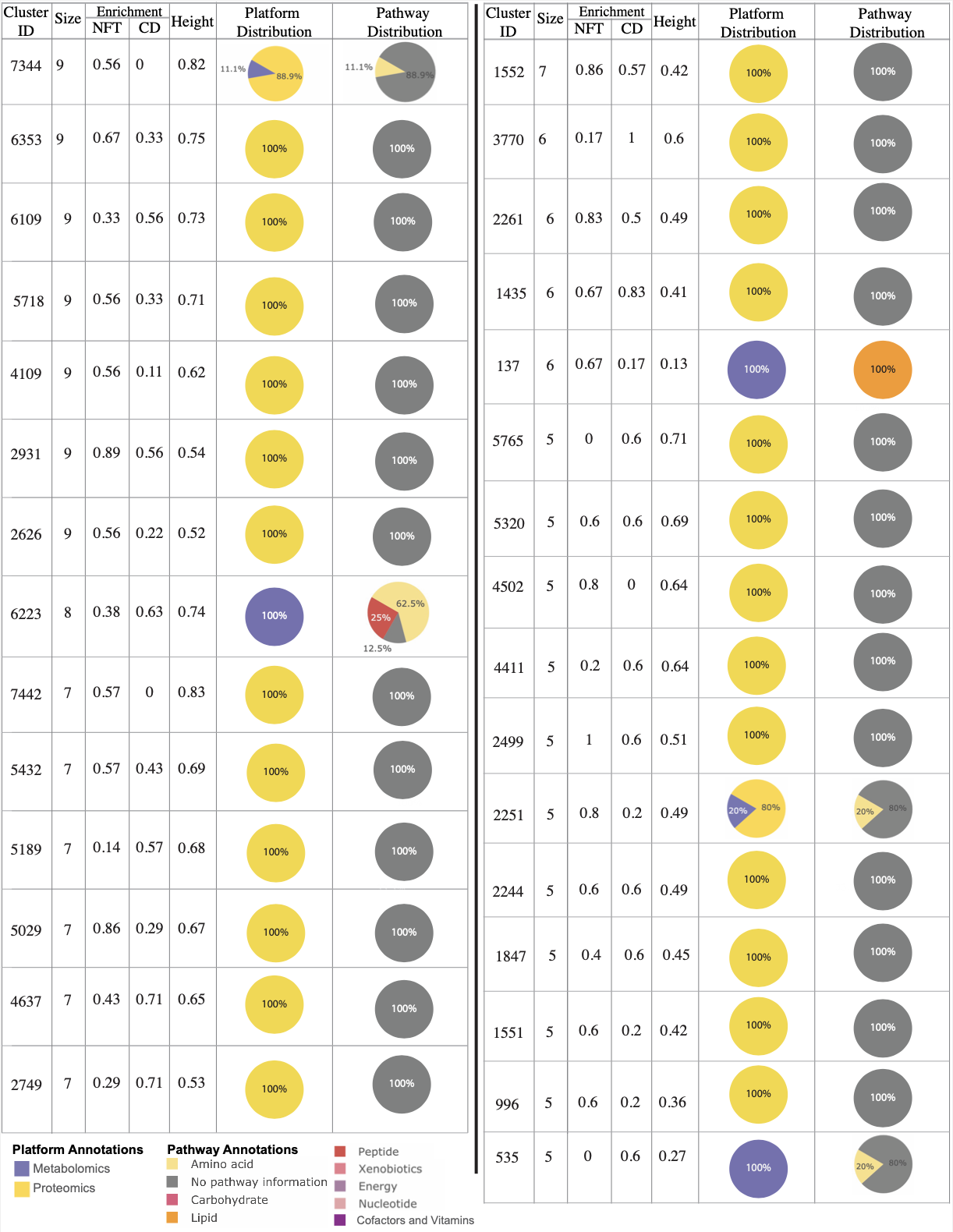

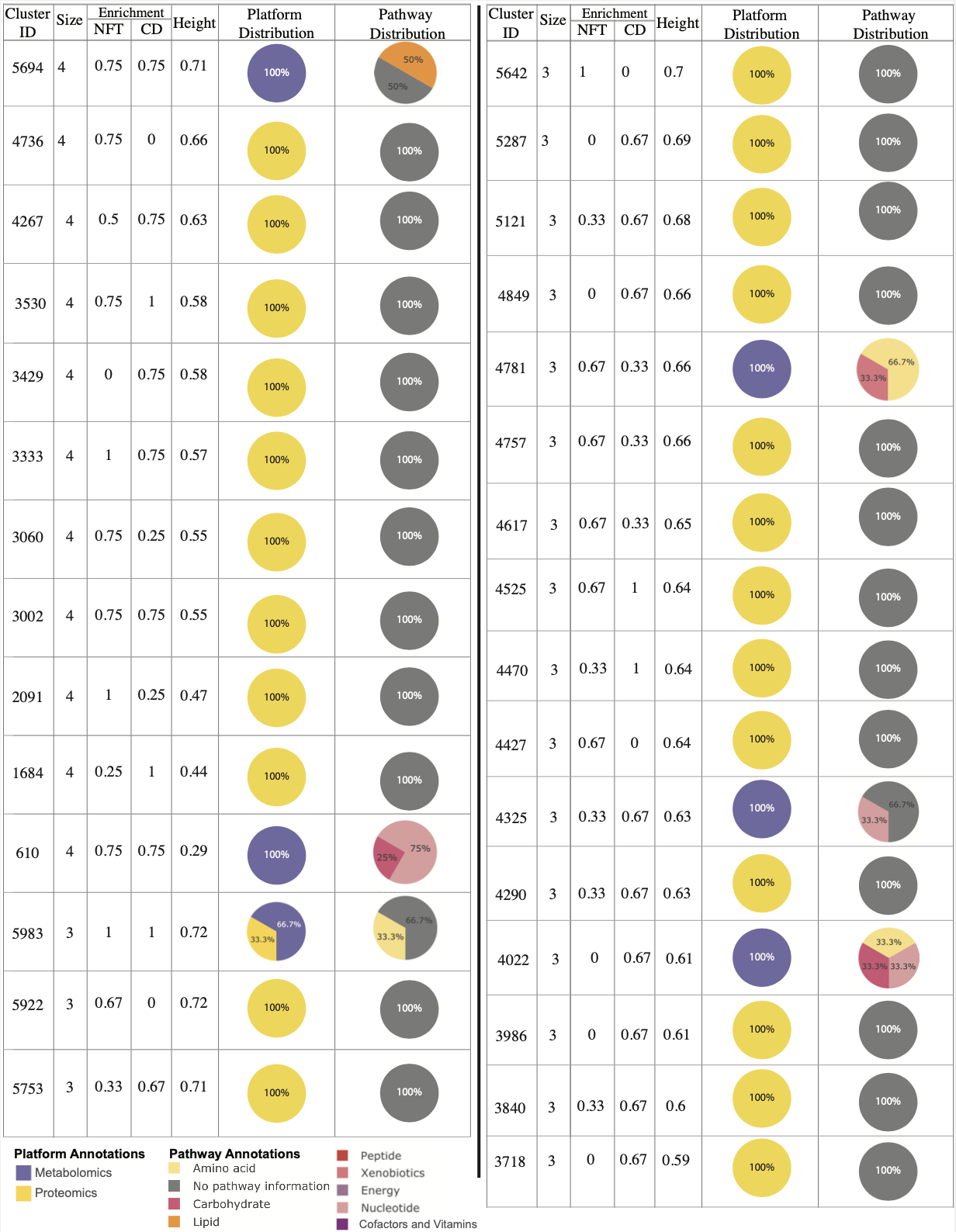

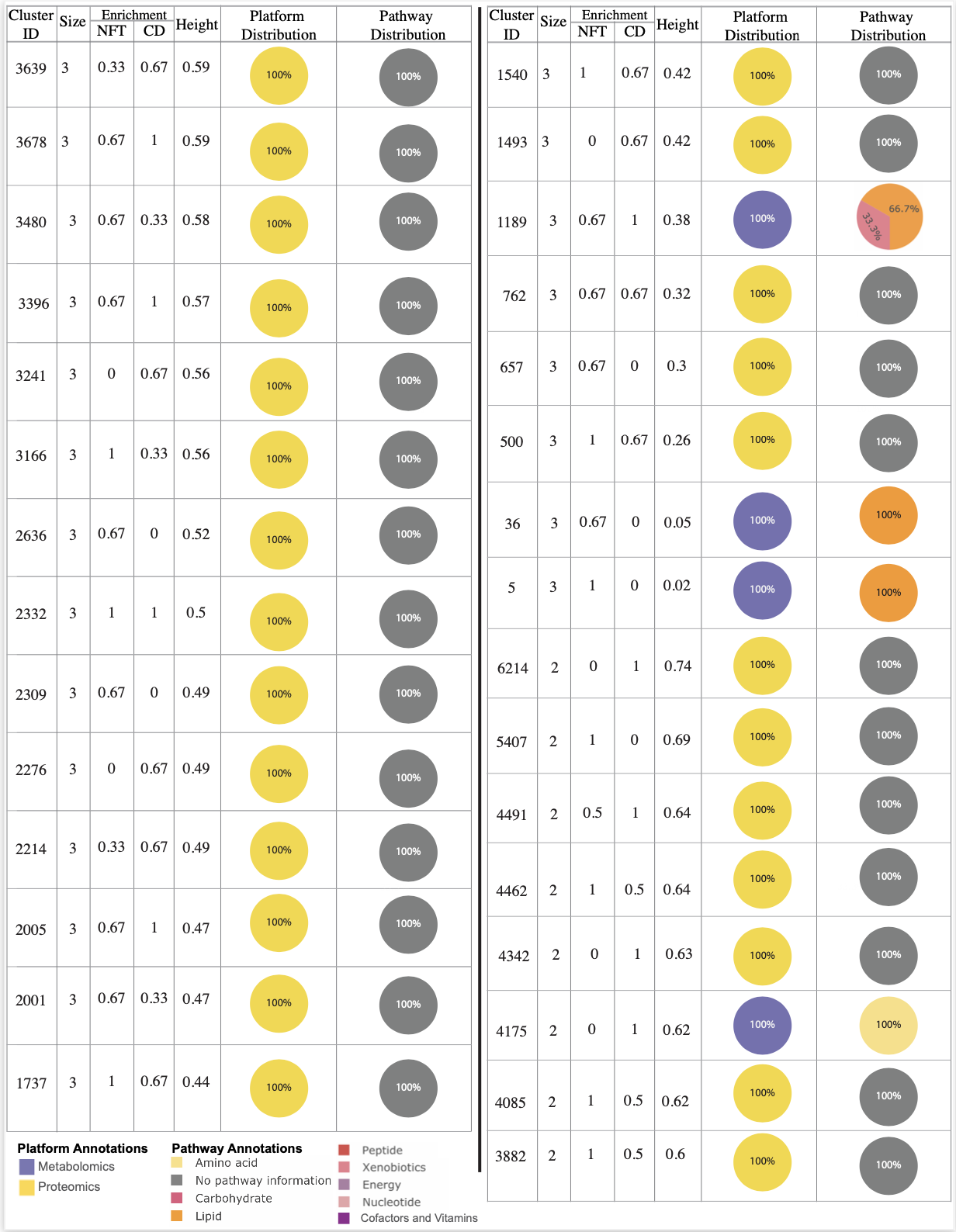

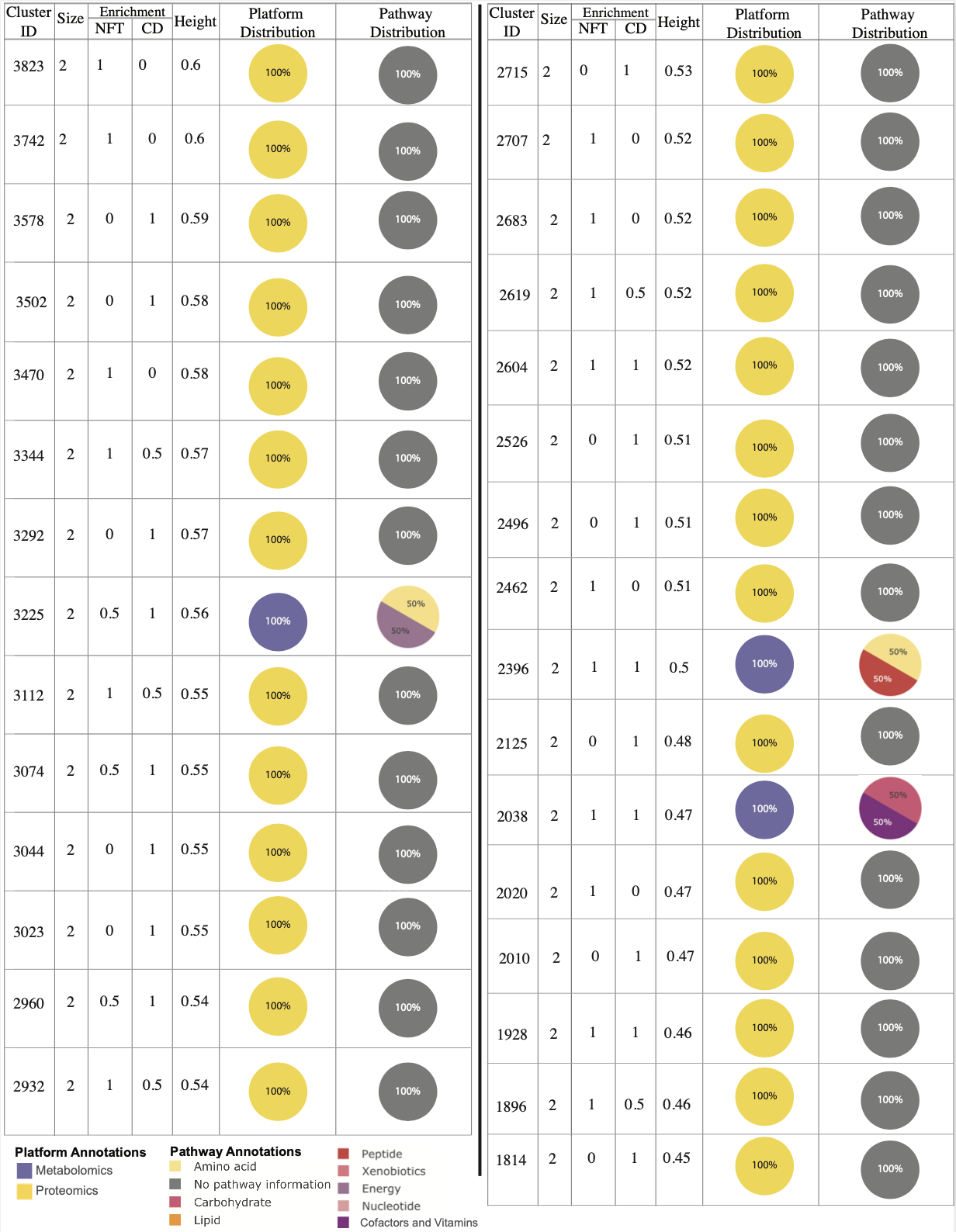

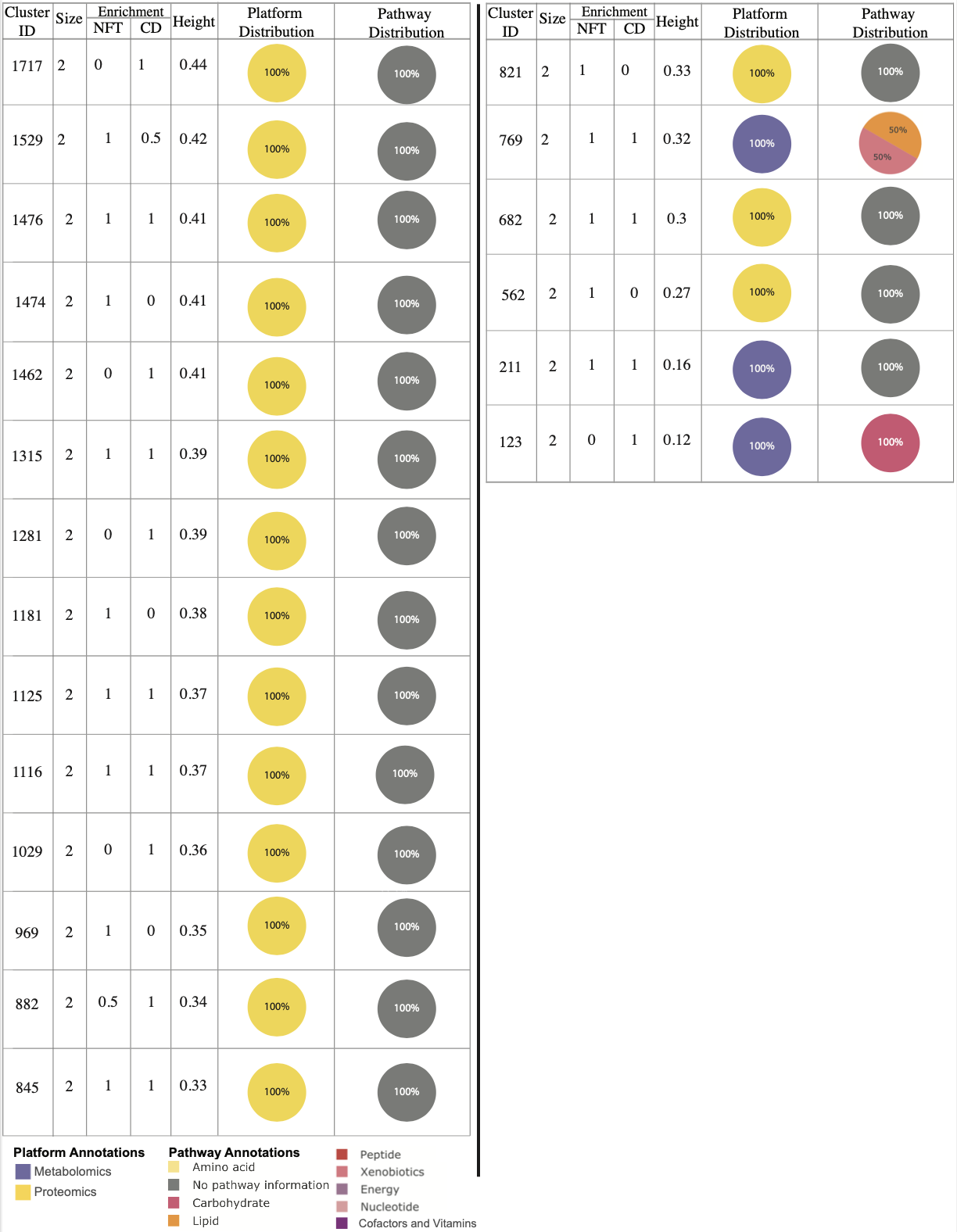

**Supplementary Figure 3.** Table of ROS/MAP module information. Listed are the modules’ ID (for cross referencing with **Supplementary Table 2**), size (number of molecules), Neurofibrillary Tangles (NFT) and Cognitive Decline (CD) phenotype enrichment (proportion of significant phenotype-associated molecules), height on the hierarchy, and distributions of the members’ datasets and pathways (metabolomics platform only).

### Supplementary Figure 4. QMDiab analysis with WGCNA clustering method

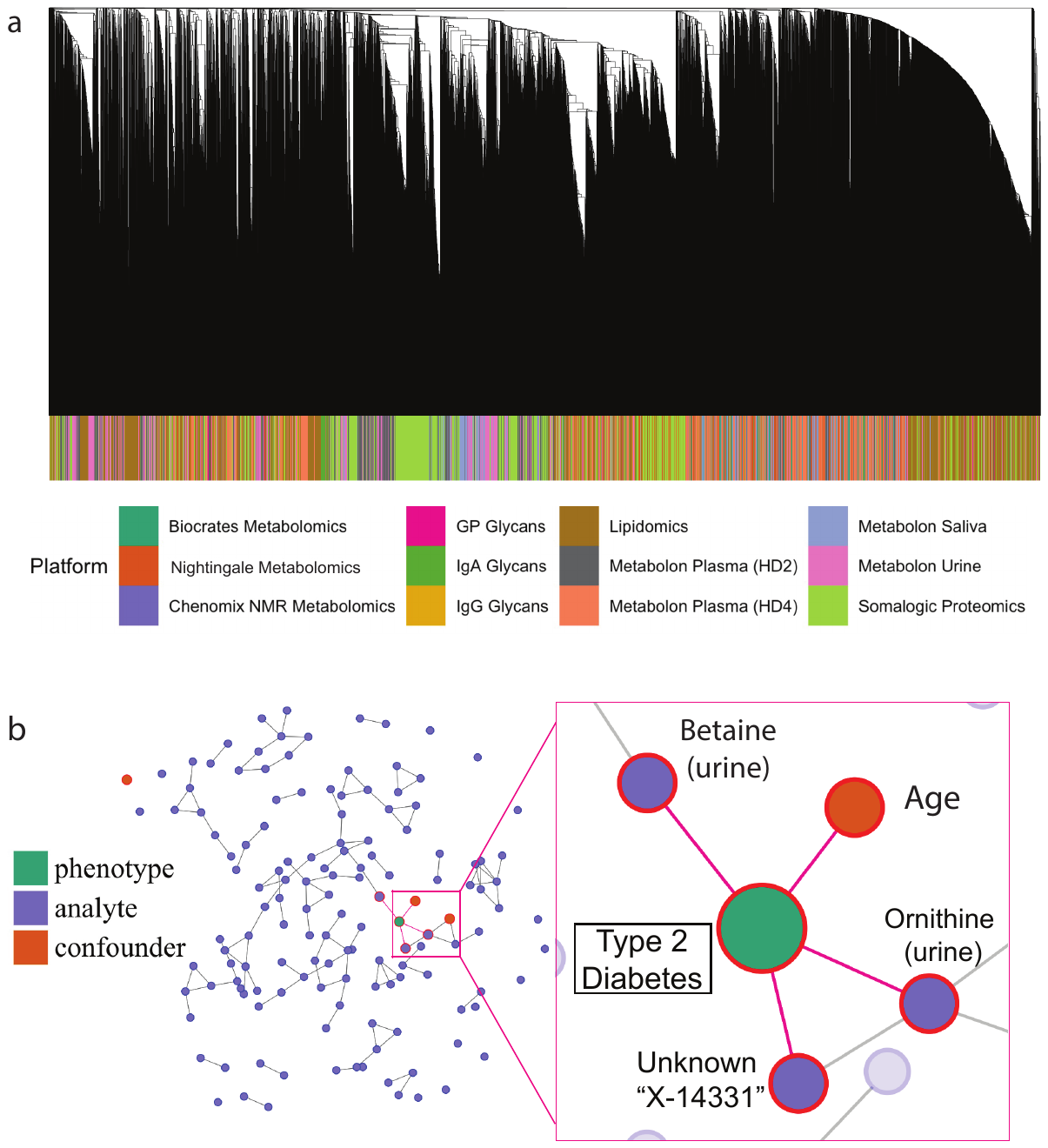

**Supplementary Figure 4. a)** Platform distribution in the WGCNA hierarchical structure on the QMDiab dataset. Strong intra-platform correlations can be seen for proteomics (green) and to a lesser extent for urine metabolomics (pink). **b**) Mixed graphical model of the 153 molecules in the largest cluster of the WGCNA analysis with phenotype and confounders. To the right, a zoomed in view of nodes with edges to the Type 2 Diabetes phenotype which include ornithine and betaine in urine along with the confounder age and one unknown molecule. Both ornithine in urine and that specific unknown were found with AutoFocus’ original hierarchical clustering method.

To investigate how a clustering method can affect AutoFocus cluster analysis, the QMDiab dataset was rerun using WGCNA’s TOM-based hierarchical structure^4^. This structure was similarly susceptible to the problem of platform bias within the resulting hierarchy; while the TOM-based hierarchical clustering more substantially distributed the lipidomics data with the other platforms as compared to correlation-based clustering, the proteomics data was largely segregated, as large regions of the tree contain only proteins (**Supplementary Figure 4a**).

The T2D enrichment step identified 13 modules within the TOM hierarchy, including a large, multi-omic cluster of 153 molecules that contained many of the same molecules as the energy metabolism module found with the correlation-based hierarchy. In contrast to the results in the main manuscript, no bone degradation proteins were identified in this cluster, potentially due to the proteomics clustering bias mentioned above. This difference in module composition also changed the MGM driver analysis, as all the drivers identified in the 153-molecule module were from urine platforms (**Supplementary Figure 4b**). Despite this, both ornithine and one unknown metabolite (labeled “X-14331”) were maintained across these two analyses.

It is worth noting that when the hierarchy is created using a TOM-based distance matrix and average linkage as is used in the original WGCNA analysis^4^, the resulting tree can have a very high maximum depth (the maximum number of internal nodes between the tree’s root and a leaf). In the instance of large datasets such as QMDiab, a high maximum depth hinders tree visualization due to memory limitations. As such, results presented here were explored without the aid of AutoFocus’ visual interface.

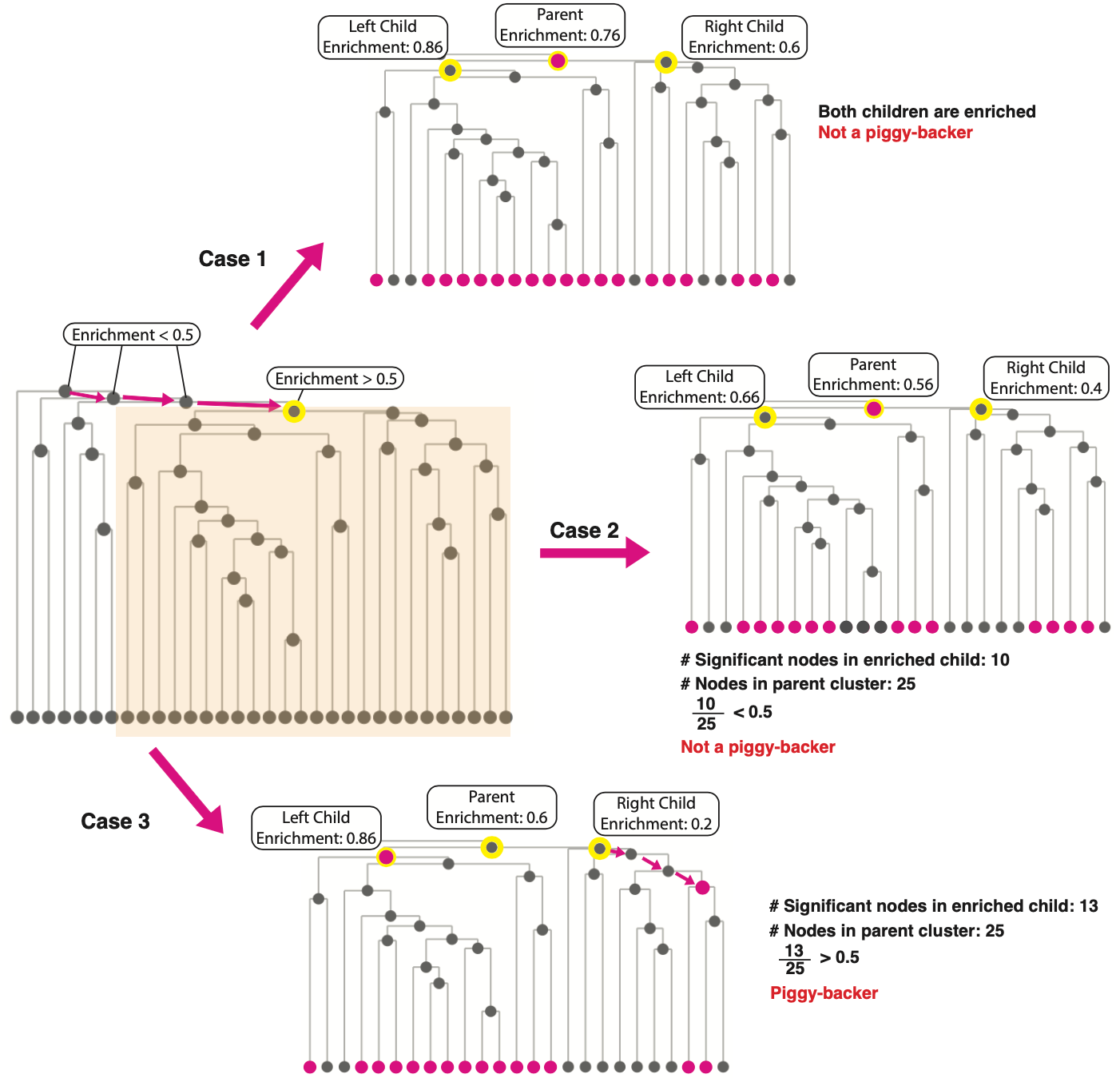

### **Supplementary Figure 5 - Enrichment Peak Identification and “Piggy-backers”**

**Supplementary Figure 5.** Example visualization of “piggy-backer” filtration process through example scenarios.

When traversing the hierarchical structure from root to leaves to identify enrichment peaks, we run into the issue of “piggy-backers”, which are defined as peaks that reach the enrichment only due to one child reaching the enrichment threshold, and the joining of the two children dilutes the signal (reduces the fraction of significant molecules in the cluster). To filter out these piggy-backers, we implement the following workflow:

Once an internal node in the hierarchical structure is found that surpasses our threshold (in this example, the threshold will be set as 0.5), we initially label it as a peak (**Supplementary Figure 5, left**). This node will be referred to as the “Parent”. It consists of an internal node descendant to the left (“Left Child”) and to the right (“Right Child”). This will result in three possible situations:

1. Both the Left Child and the Right Child also surpass the enrichment threshold. In this case, both children are enriched for signal, meaning neither is diluting the signal and therefore the Parent is a true peak and not a piggy-backer (**Supplementary Figure 5**, Case 1).
2. Only one Child meets the enrichment threshold. Without loss of generality, let the Left Child have a higher enrichment than the Right Child. In this case:
   a. If the number of significant children in the Left Child is not enough for the Parent cluster to surpass the threshold (i.e. # significant children in Left Child cluster/ # nodes in Parent Cluster < 0.5), this means that the significant nodes in the Right Child were contributing to rather than diluting the Parent Cluster’s enrichment, and therefore the Parent is a true peak and not a piggy-backer **(Supplementary Figure 5**, Case 2).
   b. If the number of significant children in the Left Child *is* enough for the Parent cluster to surpass the threshold (i.e., # significant children in Left Child cluster/ # nodes in Parent Cluster > 0.5), this means that the Parent cluster could be enriched only due to the Left Child without the contribution of the Right Child. The Parent is then deemed a piggy-backer and the Left Child is labeled as a peak. The Left Child will then undergo the same piggy-backer identification process as the Parent, and the tree will continue to be traversed down the Right Child (**Supplementary Figure 5**, Case 3).
